# Movement imitation via an abstract trajectory representation in dorsal premotor cortex

**DOI:** 10.1101/294207

**Authors:** Aaron L. Wong, Steven A. Jax, Louisa L. Smith, Laurel J. Buxbaum, John W. Krakauer

**Author notes:** Corresponding author: 50 Township Line Rd., Elkins Park, PA 19027. Co-first authors (Aaron L. Wong, Steven A. Jax). The authors declare no competing financial interests.

## Abstract

Humans are particularly good at copying novel and meaningless gestures. The mechanistic and anatomical basis for this specialized imitation ability remains largely unknown. One idea is that imitation occurs by matching body configurations. Here we propose an alternative route to imitation that depends on a body-independent representation of the trajectory path of the end-effector. We studied a group of patients with strokes in the left frontoparietal cortices. We found that they were equally impaired at imitating movement trajectories using the ipsilesional limb (i.e., the non-paretic side) that were cued either by an actor using their whole arm or just by a cursor, suggesting that body configuration is not always critical for imitation and that a representation of abstract trajectory shape may suffice. In addition, imitation ability was uncorrelated to the ability to identify the trajectory shape, suggesting a dissociation between producing trajectory shapes and perceiving their paths. Finally, a lesion-symptom mapping analysis found that imitation deficits were associated with lesions in left dorsal premotor but not parietal cortex. Together, these findings suggest a novel body-independent route to imitation that relies on the ability to plan abstract movement trajectories within dorsal premotor cortex.

**Significance Statement:** The ability to imitate is critical for rapidly learning to produce new gestures and actions, but how the brain translates observed movements into motor commands is poorly understood. Examining the ability of patients with strokes affecting the left hemisphere revealed that meaningless gestures can be imitated by succinctly representing only the motion of the hand in space, rather than the posture of the entire arm. Moreover, performance deficits correlated with lesions in dorsal premotor cortex, an area not previously associated with impaired imitation of arm postures. These findings thus describe a novel route to imitation that may also be impaired in some patients with apraxia.

## INTRODUCTION

Imitation is critical for communicating through gesture, using tools, and acquiring new motor skills. The study of imitation has revealed that humans, unlike all other animals including non-human primates, are uniquely adept at copying meaningless gestures that have no evident goal or desired outcome other than the action itself (Whiten et al., 2004; Subiaul, 2016). The processes that comprise this ability remain poorly understood.

Current theories of imitation largely stem from research into ideomotor apraxia. In this intriguing condition, caused primarily by a left hemisphere lesion, patients have relatively normal reaching and grasping (Buxbaum et al., 2005; Ietswaart et al., 2006) but cannot imitate (Buxbaum et al., 2014) even when using their non-paretic ipsilateral arm. Thus apraxia is thought to be a higher-order motor disorder. This prior work has suggested that humans imitate by moving limb segments to match the body configuration of the actor (Goldenberg, 2009; Goldenberg, 2013). This theory, however, has primarily arisen from studying the imitation of static postures (Goldenberg, 1999) rather than dynamic gestures.

Imitating moving gestures becomes computationally challenging if a description of the changing body configuration is required, as it would require keeping track of the motion of each limb segment. A simpler approach would be to represent a gesture as the path traced by the end-effector (i.e., the movement trajectory). For example, when writing the letter “S” it is sufficient to specify only the sinuous shape along which the end-effector moves, rather than a sequence of arm configurations. Indeed, such abstract trajectory-path representations can explain kinematic invariance in handwriting across effectors (Wright, 1990; Rijntjes et al., 1999) or size scales (van Galen and Teulings, 1983; Wing, 2000; Kadmon Harpaz et al., 2014). We have recently demonstrated that a trajectory-path representation is also used to plan reaches around obstacles (Wong et al., 2016). Moreover, this may explain how non-primates such as dolphins imitate human actions despite lacking multi-segmented arms (Harley et al., 1998; Xitco Jr et al., 1998). We therefore hypothesize a second route to imitation: one that does not specify body configurations, but is instead trajectory-based and body-independent.

If there are two routes to imitation, it is reasonable to ask whether these routes are functionally and anatomically dissociable. Current evidence suggests that imitation is associated with neural activity in the left inferior parietal and premotor cortices (i.e., Brodmann areas 39 and 40, and 6) (Molenberghs et al., 2009; Caspers et al., 2010; Buxbaum et al., 2014). However, there is likely a caudal-rostral perceptual-motor gradient in the brain as well as a division of the dorsal stream into dorso-dorsal and ventro-dorsal routes (Rizzolatti and Matelli, 2003; Binkofski and Buxbaum, 2013), suggesting these regions may serve different roles in imitation. Interestingly, neural activity in dorsal premotor cortex (PMd) has been associated with planning movement paths to a target (Hocherman and Wise, 1990, 1991; Pearce and Moran, 2012); lesions in PMd impair planning reaches around obstacles, resulting in attempted reaches directly toward the target through the barrier (Moll and Kuypers, 1977). Thus, we hypothesized that if there is a route to imitation involving the representation of a movement trajectory, the neural substrates of such a representation are maintained in PMd and not inferior parietal lobe (IPL).

To address our behavioral and anatomical hypotheses, we required neurotypical controls and patients with left hemisphere strokes to imitate meaningless movement trajectories. Imitation was tested using the left hand, corresponding in patients to the ipsilateral (unaffected) limb. Critically, participants were cued by watching either a cursor (providing no body-configuration information) or an actor. We reasoned that if patients were equally impaired at imitating both stimuli, this would support a body-independent route to imitation. Patients had heterogeneous lesion locations that, as a group, included PMd and IPL, with the anticipation that patients would exhibit a range of imitation abilities depending on lesion size and location. This enabled us to test where in the brain movement-trajectory paths may be computed.

## METHODS

Twenty-two right-handed chronic stroke survivors with lesions confined to the left hemisphere (8 female) and thirteen right-handed neurologically intact controls (7 female) completed the experiment. One patient was subsequently removed from the study prior to analysis due to impaired visual abilities. The remaining group of patients (Table 1) was marginally younger than the control group (patients: 57.4 ± 10.1 years, range 35-80 years; controls: 64.2 ± 11.1 years, range 44-80 years; p = 0.086), thus any detrimental effects of aging on performance would be reflected more heavily in the control group than the patient group. All participants provided written consent to participate in the study in accordance with the guidelines of the Einstein Healthcare Network Institutional Review Board and were compensated for their participation.

**Table 1.**
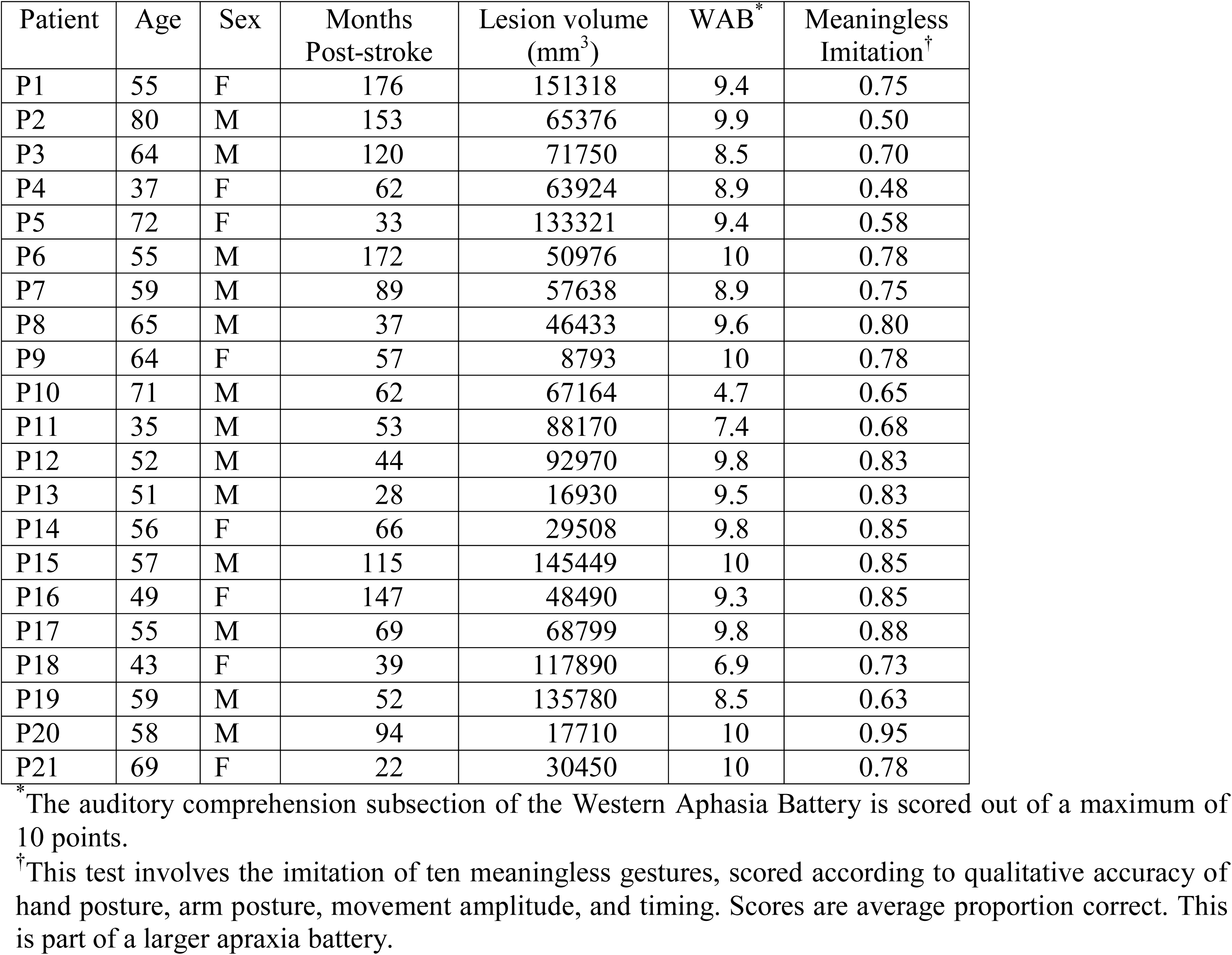
List of patient participants with left-hemisphere strokes

All participants completed two tasks as part of this experiment: the Production task and the Perception task (details below). All tasks involved a set of trajectory shapes that were chosen such that the resulting gestures would be relatively arbitrary and did not have a commonly associated meaning. (Note, although it is possible to use a heuristic to link these gestures to meanings, such a strategy could be applied to any potential gesture selected for this task; hence our selection criterion was simply that the entire trajectory of the movement, considering both spatial and temporal aspects, had no obvious meaning). Each task consisted of two trial phases. The first phase was a practice block in which participants were provided with instructions on how to complete the task, were shown how to perform the task, and then allowed to rehearse three example trials in which they were given feedback regarding their performance. Following this, participants completed a test phase during which they received no further reminders of task instructions or feedback regarding their accuracy in performing the task. Participants received different stimuli during the practice and test blocks.

### Trajectory production task

In the Production task, participants executed two dimensional movements by sliding their fingertip along the surface of a vertically mounted sheet of clear Plexiglass. The left arm was used for all participants so that the stroke group would produce movements with the ipsilateral limb. Stimulus trajectories varied in size, but all were contained within a 38 cm x 56 cm area at the center of the board. Participants sat at a distance that allowed them to comfortably perform movements, approximately 50 cm from the Plexiglass with their hand resting in their lap (i.e., the “home” position). Instructions about the movements to perform were presented via a laptop computer placed approximately 30 cm behind the Plexiglass. Each video clip, which ranged in length from 4.5 to 6.4 seconds, was played twice with a one second delay between clips. Following the second video, there was a one second delay before a tone sounded to inform participants they should begin their movement. The movement consisted of reaching out and upward to contact the Plexiglass surface, generating the required trajectory in the vertical plane by sliding the hand along the Plexiglass, and then returning the hand to the home position in their lap. Participants were given the instructions to “use you left finger to trace the same path the dot or person made. Make sure your path looks exactly like the video’s. It’s very important that you be as precise as possible.” If participants began to move before the end of the video, they were reminded to wait for the tone. The position of each participant’s left index finger was continuously recorded using an Ascension TrakSTAR motion tracking system (Ascension Technology, Shelburne, VT) at 150 Hz.

Participants produced 16 trajectories (Figure 2) grouped into two presentation conditions. We manipulated task difficulty by providing a differing number of visible landmarks (2 cm diameter circular dots placed on the Plexiglass) under the hypothesis that landmarks could potentially be used as reference points to subdivide a complex action into simpler point-to-point reaches, thus reducing the trajectory planning requirements. Stimuli were thus cued with reference to either a single central landmark or with four additional landmarks bounding a rectangular workspace. However, in general we found that varying the number of landmarks did not significantly impact performance; thus, while we include a landmark factor in our statistical models for completeness (see Quantification and Statistical Analysis below), for conciseness we do not report the outcomes of statistical tests of this factor in the Results.

More importantly, each trajectory was reproduced under two levels of stimulus type (body or cursor)^1^. As described in the Introduction, this stimulus-type manipulation was included to test whether trajectory imitation is dependent upon the ability to observe and represent the relative positions of limb segments during the action, or if trajectory imitation instead can be performed more abstractly by replicating only the kinematics of the end-effector. Video stimuli for the body condition consisted of a seated actor producing trajectories on a Plexiglass board identical to the one in front of the participant (i.e., with the same number of landmarks). The actor was shown facing the participant and moving his right arm so that the experimenter’s arm mirrored the participant’s left arm. Participants were trained to imitate the mirrored version of the actor; no mirror-inversion mistakes were observed for either the patients or the controls during this task. Video stimuli for the cursor condition consisted of a single black cursor moving over a white background, with landmarks visible as appropriate. Cursor trajectories were created from motion tracking data collected during filming of the body stimuli, resulting in identical end-effector movement trajectories for the two stimulus types. Participants completed two blocks of trials; the two levels of the landmark factor (1 or 5) were manipulated between-blocks, while the two levels of the stimulus type factor (body or cursor) were intermixed within blocks.

As part of this Production task, we also examined the ability of patients and controls to imitate simple point-to-point reaches by evaluating the ability to imitate the initial segment of one of the more complex movement trajectories (see below for details). Analysis of point-to-point movements allows us to control for whether observed imitation deficits could be attributed to a lower-level deficit in movement execution.

### Trajectory perception task

A deficit in imitating trajectories could be attributed to one of two possible causes: impairment in the ability to plan the necessary trajectory shape, or difficulty with perceiving and/or recalling the required shape (i.e., a perceptual and/or working memory problem). To distinguish between these possibilities, participants were additionally required to complete a Perception task. In this task, participants were exposed to the identical video stimuli as in the Production task, but were asked to simply make a decision about the path they had observed rather than producing the actual movement. That is, stimuli consisted of two presentations of the same videos used in the Production task followed by a one second delay before the presentation of a static image of a trajectory shape. Participants were instructed to indicate “if the path in the picture is the same as the movement made in the video.” The shape either matched the exact trajectory produced in the video or was a mirror image of the trajectory that was reflected about the vertical or horizontal axis. The shape remained on the screen until participants indicated a match or non-match by pressing one of two buttons on a computer keyboard with their left hand. Task accuracy and reaction time (RT) were measured, although due to technical difficulties RT data at the trial level were unavailable for one control and two stroke participants, and thus these individuals could not be included in analyses involving mixed-effects models (see Quantification and Statistical Analysis below).

As with the Production task, participants in the Perception task completed two blocks of trials; the two levels of the landmark factor (1 or 5) were manipulated between-blocks, and the two levels of the stimulus type factor (body or cursor) were intermixed within blocks. The order of the Production and Perception tasks were counterbalanced across participants.

Participants were also required to complete a task intended as an independent test of visual working memory ability with stimuli that were not trajectories. In this Memory control task, participants were shown one of eight images chosen from a set of abstract line drawings (Petrides and Milner, 1982). Each image was presented twice for 4 seconds with a one second delay in between; following a one second pause, a second stimulus was displayed which was either identical or a rotated version of the first image, and participants were required to report whether the two images were the same or different using a keypress with their left hand. Thus, the structure of this task exactly mirrored that of the Perception task, except the initial stimuli were static pictures instead of videos. Task accuracy and RT were measured using the same method as described below for the Perception task. Although average task accuracy did not differ between patients and controls (t-test, *t*(32) = 0.51, *p* = 0.61), average RT was different between groups (t-test, *t*(32) = −2.67, *p* = 0.01) and so was introduced as a covariate when appropriate in the analyses involving RT below. The Production task data as well as the Region of Interest (ROI) Lesion-Symptom Mapping analyses were also re-examined with inclusion of average RT from the Memory task analysis as a covariate to control for any potential visual working memory deficits in the patient group, but since inclusion of this additional term did not change the significance of any findings, we do not report the results of those models here.

### Stroke group auditory comprehension

Patients with strokes completed the auditory comprehension subsection of the Western Aphasia Battery (Kertesz, 1982), which quantifies capacity to understand verbal instructions. Scores on the WAB were high (9.06 ± 1.32 out of 10 possible points). Thus, we felt confident that any group differences in performance could not be explained by differences in instruction comprehension.

## Statistical Analysis

### Production Task

Response trajectories were recorded and analyzed offline using custom software written in Matlab (The MathWorks, Natick, MA). Fingertip velocity was calculated by smoothing the recorded fingertip position using a second-order Savitzky-Golay filter with a frame size of 19 samples, then taking the derivative. Movement start and end were identified as the time when the fore/aft reach velocity became minimal (< 0.03 m/s), reflecting the time when the participants contacted the Plexiglass surface and their primary axis of motion was along this surface. Each start and end time was verified by visual inspection to remove erroneous points such as if a participant lifted their hand away from the Plexiglass in the middle of the trajectory. All motion produced between the identified start and end times was labeled as the complete imitated movement generated by the participant. The three-dimensional data returned by the motion tracking system were projected into the plane of the Plexiglass surface to account for any small angular mismatch between the Plexiglass (on which the reference landmarks were positioned) and the vertical plane of the tracker axes. Movements were smoothed and time-normalized using a b-spline fit with 200 points. Finally, the shape of the generated response was compared to the ideal template (the normalized and smoothed trajectory produced by the actor in the stimulus video) by performing a Procrustes distance analysis (Goodall, 1991). The Procrustes distance analysis estimates the best translation, rotation, and scaling parameters that match the sample reach to the template, and returns a dissimilarity measure for the resulting transformed response trajectory shape (normalized between 0 and 1, with 0 being identical and 1 being completely dissimilar).

Data were analyzed using mixed-effects models including fixed effects of Group (patient or control), Stimulus type (body-cued or cursor-cued), Landmark (1-dot or 5-dot), Task order (Production first or Perception first), and random effects of Subject and Stimulus shape. Models were fit using the *glmmTMB* package in R (Magnusson et al., 2017) to improve convergence and to allow for the use of a logarithmic link function in the model to address the non-Gaussian (bounded between 0 and 1) nature of the Procrustes distance metric. Significant main effects were evaluated using likelihood ratio tests comparing a model with the effect of interest against a model without the effect of interest (which approximately follows a chi-square distribution). Step-down Bonferroni-Holm corrections were applied to address multiple comparisons across likelihood ratio tests.

### Trajectory production control: point-to-point reaches

Although patients were tested on their non-paretic side, we wanted to confirm that errors made during trajectory production could not be explained by a low-level motor execution deficit. Thus, we isolated point-to-point reaches from the initial movement segment of one of the 5-landmark stimuli for both the body and cursor conditions. This particular stimulus was chosen because it afforded a reasonably long initial movement segment that started and ended on visible landmarks, thus providing clearly identified movement start and end positions that defined the desired movement. The movement segment of interest was identified using a velocity threshold (tangential velocity > 0.05 m/s), and was verified by visual inspection to address cases where participants did not completely pause between segments. To evaluate the quality of reproducing the point-to-point movement, we calculated the maximal reach deviation as the greatest absolute perpendicular deviation of the finger from the straight line between the initial and final fingertip position of the point-to-point reach. We also calculated the initial reach direction of the velocity vector 100 ms after movement onset, and the initial direction error as the signed difference between the initial reach direction of the participant and the ideal (actor) reach direction.

Data were analyzed with mixed-effects models with main effects of Group and Stimulus type (body-cued or cursor-cued), and random effect of Subject regressing against maximal reach deviation from a straight line or initial direction error. Models were fit using the *lme4* package in R (Bates et al., 2015). Significant effects were evaluated using likelihood ratio tests. Here and elsewhere, error bars reflect S.E.M.

### Perception Task

Reaction time (RT) was measured as the time between the onset of the static trajectory and initiation of a button-press response (reporting “same” or “different”). RT data were trimmed by first removing the RTs on incorrect trials (since incorrect responses might arise from non-task-related causes such as lack of attention on a particular trial), then removing all RT outliers falling more than 2.5 standard deviations of the mean. Accuracy and RT were analyzed with mixed-effects models with main effects of Group, Stimulus type (body-cued or cursor-cued), Landmark (1-dot or 5-dot), and Task order (Production first or Perception first), and random effects of Subject and Stimulus shape. In the case of RT, average RT performance in the Memory task was also included as a fixed effect to control for individual differences associated with working memory. For all behavioral tests, the logarithm of the RT (logRT) was used to reduce outliers and address the long-tail distribution, as it is well recognized that the logRT distribution is approximately Gaussian (e.g., (Carpenter, 1981)). Models were fit using the *glmmTMB* package in R (Magnusson et al., 2017), using a Gaussian link function in the case of RT or a binomial family in the case of Accuracy. Significant main effects were evaluated using likelihood ratio tests comparing a model with the effect of interest against a model without the effect of interest; step-down Bonferroni-Holm corrections were applied to address multiple comparisons.

Correlations in performance between the Perception and Production tasks were tested by examining the relationship between mean Procrustes distance and mean accuracy or mean logRT, with means taken within each subcondition (i.e., for each unique Stimulus type and Landmark). Means were used in this analysis to allow for calculation of a Percentage Accuracy score in the Perception task while preserving potential differences arising from the effects of stimulus type or landmark. Mixed-effects models also included fixed effects of Group (patient, control), Stimulus type (body-cued or cursor-cued), and Landmark (1-dot or 5-dot), and random effects of Subject. Models were fit using the *lme4* package in R (Magnusson et al., 2017). Significant main effects of the effect of interest (i.e., Accuracy or RT) were evaluated using likelihood ratio tests comparing a model with the effect of interest against a model without the effect of interest.

### Lesion location identification

T1-weighted MRI brain scans (15 patients) or CT scans without contrast if MRI was contraindicated (6 patients) were used to identify lesion locations; data from all 21 patients were included in the analyses listed below. Lesions were segmented under the supervision of a neurologist (H. Branch Coslett) who was blind to all behavioral data. Details of the lesion location identification procedure can be found in Kalénine et al. (2010). Individual lesion locations for all stroke participants are shown in Figure 1. Total lesion volume for each individual patient was calculated (71,850 ± 34,488 mm^3^) and used in some analyses as a control for overall stroke severity (see below).

**Figure 1.**
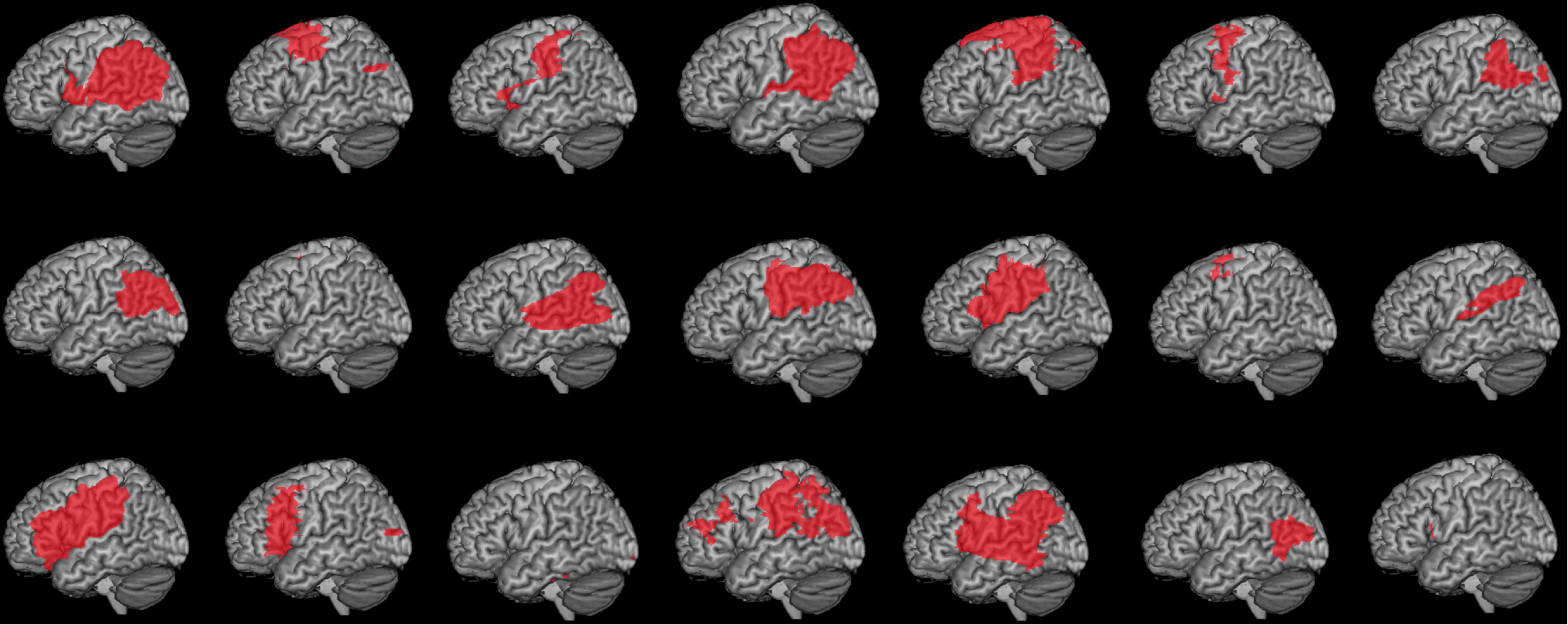
Cortical view of lesion maps for individual patients. Patients were selected to have a range of lesions spanning the left fronto-temporo-parietal cortex.

### Multivariate lesion-symptom mapping

Multivariate lesion-symptom mapping was conducted using support-vector regression lesion-symptom mapping (SVR-LSM) (Zhang et al., 2014), a machine learning technique for multiple regression that trains a model to predict behavioral scores using the full lesion maps of all participants. SVR-LSM is an improvement over earlier techniques such as voxel-based lesion-symptom mapping (VLSM) (Bates et al., 2003), which perform all analyses at the individual voxel level and then aggregate across voxels to create a full-brain map. By predicting behavioral scores using all voxels simultaneously, SVR-LSM has the capability to avoid vasculature-related biases that can systematically distort lesion-symptom maps (Mah et al., 2014). SVR-LSM analyses were performed using software written by Zhang and colleagues (Zhang et al., 2014; https://cfn.upenn.edu/∼zewang/) and modified by DeMarco and colleagues (DeMarco and Turkeltaub, 2018). As with the behavioral analyses, mean RT during the Memory Task was included as a behavioral covariate in the SVR-LSM analysis to control for any differences in working memory across patients.

Statistical significance was determined using a voxel-wise threshold. First, a beta-map of the observed data was computed. This beta-map regressed out the effects of total lesion volume in the behavioral data as well as normalizing each subject’s lesion data vector to have a norm of one (dTLVC procedure, see (Zhang et al., 2014)) to avoid identifying spurious voxels whose lesion status correlates with lesion volume. Only voxels damaged in at least three participants (15% of the stroke group) were included in the analyses. Although this voxel-inclusion criterion is frequently set to 10% of the total number of patients (Kalenine et al., 2010), due to our small sample size we increased this threshold to 15% to decrease the number of spurious voxels included in the analysis. A voxel-wise p-map was then derived by comparing the observed beta-map to beta-maps from 10,000 Monte-Carlo style permutations where behavioral scores were randomly paired with lesion maps. For analyses averaging across the body and cursor conditions, significant clusters were identified using a cutoff threshold of *p* < 0.01 and removing all clusters smaller than 500 mm^3^. As a secondary SVR-LSM analysis we were also interested in visualizing the potential overlap in brain regions correlated with behavior within the body or cursor conditions separately; results for these analyses were shown at a relaxed threshold of p < 0.05.

In addition, we performed region of interest (ROI) analyses to confirm whether there was any relationship between performance in either the Production or Perception task and the percentage of lesioned voxels within three ROIs selected *a priori*: Brodmann area 6 (premotor cortex), Brodmann areas 39 (angular gyrus), and Brodmann area 40 (supramarginal gyrus). Mixed effects models were run using the *glmmTMB* package in R (Magnusson et al., 2017) and, aside from the three pre-selected ROIs, also included fixed effects of Total Lesion Volume, Stimulus type (body or cursor), and Landmark (1-dot or 5-dot). Models also included random effects of Subject and Stimulus shape. Significant main effects were evaluated using likelihood ratio tests comparing a model with the effect of interest against a model without the effect of interest, with step-down Bonferroni-Holm corrections applied to address multiple comparisons.

## RESULTS

Twenty-one patients with left-hemisphere stroke (Fig. 1) and thirteen age-matched controls were asked to imitate meaningless trajectories (Fig. 2) using their left (non-paretic, non-dominant) hand after watching the movement of either an actor (body-cued stimulus) or a cursor (body-free stimulus). Performance in this Production task was contrasted against a Perception task in which they simply had to report whether a subsequently displayed static image had the same or different shape compared to the stimulus trajectory.

**Figure 2.**
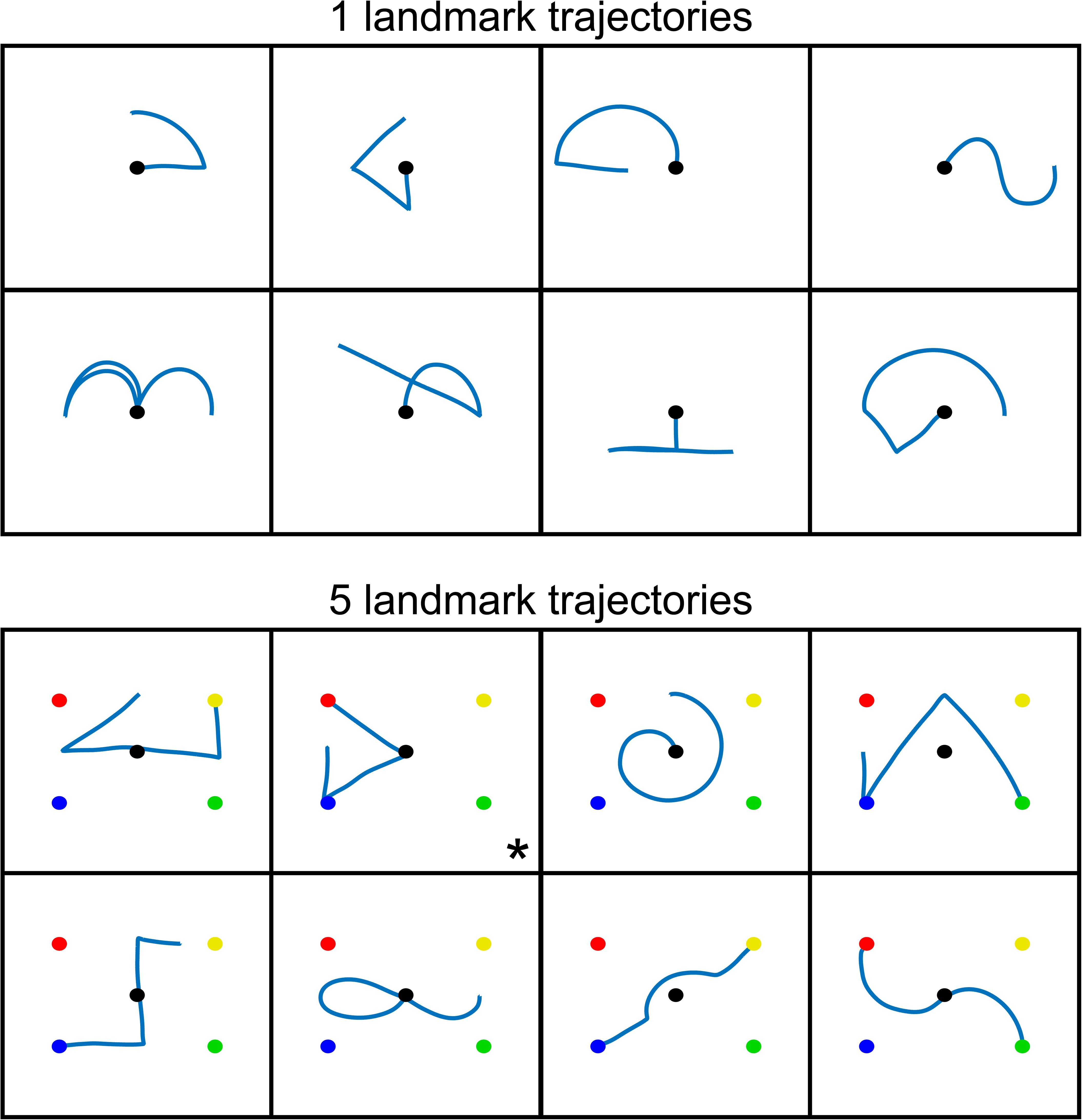
Reach trajectories employed in the Production and Perception tasks. Trajectories were cued by observing the movement of either an actor or a cursor, and were made with reference to either one (top panel) or five (bottom panel) visible landmarks. These static shapes were derived from motion tracking of the actor’s fingertip, and were displayed in the Perception task. In the one-landmark condition (top), motion always began at the central target; in the five-landmark condition (bottom), the starting location of the trajectory varied for each stimulus shape. In all cases, while participants observed a stimulus (actor or cursor) moving along these paths, a static representation of the trajectory path was never displayed as part of the stimulus. The asterisk in the second panel in the 5-landmark condition denotes the trajectory from which the initial movement segment was extracted to evaluate the quality of imitating point-to-point reaches. This trajectory was selected because it contained a single straight line at the start of the movement that started and ended on well-defined landmarks.

### Patients and controls exhibited comparable point-to-point reach kinematics

We first analyzed the imitation of point-to-point reaches to confirm that the ability to reproduce straight movement paths was unimpaired in patients. Example point-to-point movements for a representative control, a patient with dorsal premotor but without inferior parietal lobe damage (patient 6 in Figure 1), and a patient with inferior parietal lobe damage but without dorsal premotor damage (patient 1 in Figure 1) are shown in Figure 3. As can be seen by comparing the three top panels of the figure, all three participants produced similar movements. This observation was confirmed in a group-level analysis focusing on the ability to generate reaches that were straight and aimed in the correct direction. Specifically, we measured the straightness of the movement path (maximum hand-path deviation from a straight line) and the initial reach direction error (deviation of the aiming direction of the hand 100 ms after movement initiation). The patient and control groups did not differ in either the straightness of the movement path (patients: 4.28 ± 2.91 mm; controls: 3.49 ± 1.72 mm; *χ*^2^(2) = 3.26, *p* = 0.20) or the initial direction error (patients: 3.48 ± 2.96°; controls: 5.39 ± 1.79°; *χ*^2^(2) = 0.25, *p* = 0.62). This is not surprising, as patients with apraxia tend to have relatively unimpaired reaching and grasping ability in their nonparetic hand (Buxbaum et al., 2005; Ietswaart et al., 2006) unlike the difficulty in generating straight movements with the paretic limb typically observed in stroke (Dewald et al., 2001; Cirstea et al., 2003; Osu et al., 2011). Thus, patients and controls imitated point-to-point reaches with comparable accuracy, suggesting that any imitation abnormality with respect to curved trajectories cannot be attributed to an abnormality in movement execution.

**Figure 3.**
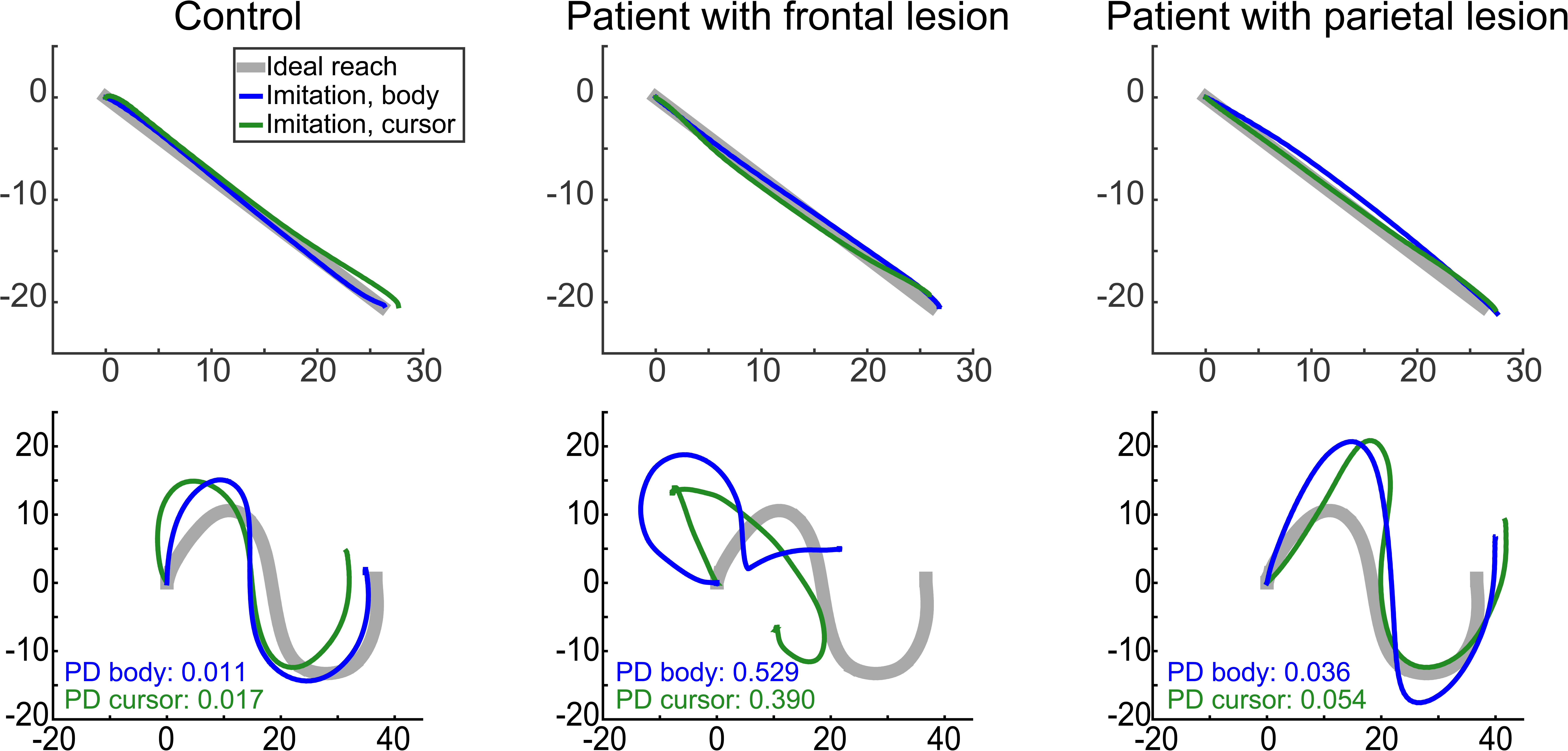
Examples of imitated trajectories from a single control participant (left column), a patient with dorsal premotor but without inferior partial damage (center column), and a patient with inferior parietal but not dorsal premotor damage (right column). Trajectories were chosen to clearly illustrate what good or bad performance looked like across patients compared to controls. In all panels, imitation of a body stimulus is shown in blue and imitation of a cursor stimulus is in green; the thick gray line reflects the ideal movement as performed by the actor. Point-to-point trajectories were comparable across participants (top row). In contrast, imitation ability for complex paths (bottom row) was worse for the patient with dorsal premotor damage than the other two participants, for both the body and cursor conditions. Imitation ability was quantified using a shape dissimilarity (Procrustes distance, PD) metric, in which larger values indicate worse replication of the actor’s movement.

### Imitation of complex trajectories was impaired in patients with left hemisphere damage

Despite being able to produce straight point-to-point trajectories, patients with left hemisphere damage exhibited deficits in the ability to reproduce the shapes of movement trajectories in the Production task (see Methods). Example trajectories are shown in the bottom panels of Figure 3. Shape imitation errors were quantified by calculating a dissimilarity score (Procrustes distance metric) (Goodall, 1991) between the movement of the actor/cursor displayed in the stimulus and the trajectory produced by the participant. Mean dissimilarity scores are shown in Figure 4 at both the group (bars) and participant (lines) level. Qualitatively, although patients exhibited a wide range of dissimilarity scores, as a group their shape dissimilarity scores were generally larger than those of the controls but similar between the body-cued or cursor-cued stimuli.

**Figure 4.**
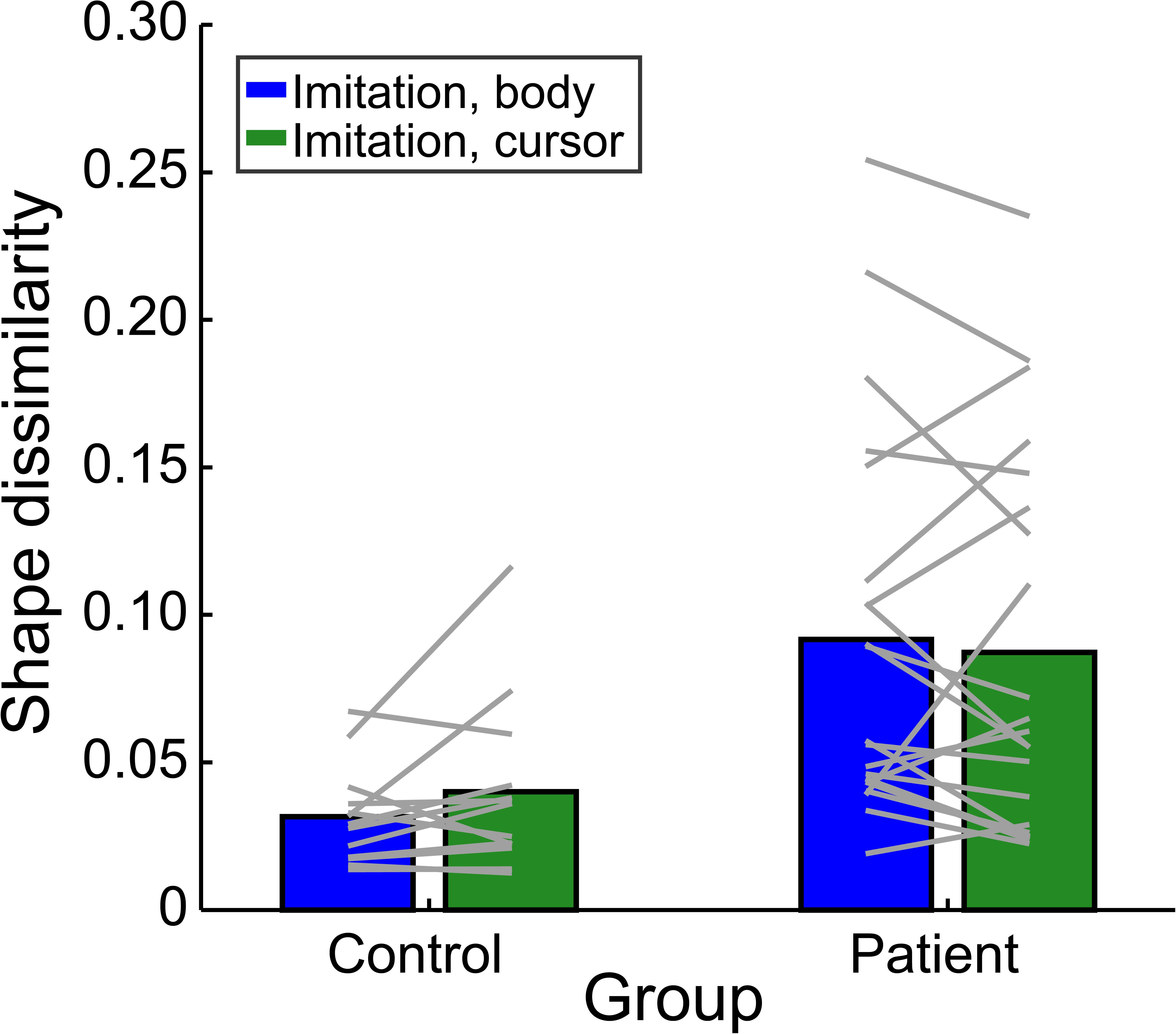
Mean shape dissimilarity (Procrustes distance) of imitated movements in the Production task. Group means (bars) and individual means (lines) are shown separately for the body-cued (blue) and cursor-cued (green) conditions, for both patients and controls.

When shape dissimilarity scores were analyzed with a mixed-effects model, there was a main effect of group (patient versus control: χ^2^(2) = 17.29, *p* = 0.0009), in which stroke participants reproduced the shape less accurately compared to controls. However, no significant main effect was observed for stimulus type (body-cued versus cursor-cued: χ^2^(1) = 0.613, *p* = 0.434), suggesting that regardless of whether or not the stimuli provided information about the relative positioning of limb segments during movement, participants performed equally well at imitating the required shape.

Consistent with the lack of a main effect of stimulus type on imitation performance, at the individual level participants who exhibited larger shape dissimilarity scores in the body condition also performed worse during the cursor condition. Regression analyses indicated that participants’ average shape dissimilarity in the body task was highly correlated with their average shape dissimilarity in the cursor task, with a slope near 1 and intercept near 0 (R^2^ = 0.78; slope = 0.902, intercept = 0.006). Similar results were obtained when considering only the stroke participants (R^2^ = 0.78; slope = 0.88, intercept = 0.015). Thus, the ability to imitate trajectory paths was not different whether participants were given information about the configuration of the entire limb or saw only a body-independent representation of the end-effector.

Although patients also moved more slowly than controls when imitating trajectories (peak velocity: controls, 0.40 ± 0.056 m/s, patients, 0.37 ± 0.056 m/s, χ^2^(2) = 9.25, *p* = 0.009), a group difference in shape dissimilarity remained even after accounting for this effect (χ^2^(2) = 7.83, *p* = 0.005) and there were no significant differences in either acceleration or jerk between the two groups (p > 0.26). Thus, it was not possible to explain these group-level differences on the basis of simpler kinematic parameters, consistent with the lack of an observed impairment at imitating straight movements as noted above. Finally, patient performance could not be explained by a nonspecific cognitive effect of lesion volume, as there was no relationship between shape dissimilarity and total lesion volume (R^2^ = 0.001, *p* = 0.857).

### Trajectory perception was also impaired in patients with left-hemisphere damage

One potential concern was that imitation deficits could arise not because of problems in generating the desired action, but as a result of deficits in perceiving or remembering the movement to be copied. Note, the findings reported here are unlikely to arise from a more general visual working memory deficit or difficulty in mental manipulation of stimuli based on an additional control task (see Methods). Hence, here we were primarily concerned with whether there was a problem in perceiving the visual motion of the stimulus and transforming it into the shape of the trajectory path. To examine this, participants were required to complete a Perception task, in which they observed a movement trajectory as in the Production task, but then simply had to make a binary choice (yes/no) as to whether a subsequently displayed static image had the same shape as the path of the viewed stimulus trajectory. Foils were static images of the correct path shape that were then reflected about the horizontal or vertical axis; as there was no significant effect of reflection direction on performance (accuracy: χ^2^(1) = 0.738, *p* = 0.39; RT: χ^2^ (1) = 1.00, *p* = 0.32), the data presented below were collapsed across reflection directions. As before, the stimuli could be cued either by an actor or a cursor.

Mean accuracy and reaction time (RT) in identifying a static shape as being the same or different from the observed movement are shown in the Figure 5A and 5B, respectively. As with the Production task, patients exhibited a wide range of performance, but were generally worse than controls (lower accuracy, higher RTs). Accuracy and RT measures were negatively correlated across participants (r^2^ = 0.25, p = 0.004), suggesting that impairments in trajectory perception generally affected both measures similarly. However, since accuracy is informative about the likelihood of correctly reporting the observed trajectory whereas RT as a continuous measure provides a sense of uncertainty about that choice, both provide complementary information about performance in this Perception task.

**Figure 5.**
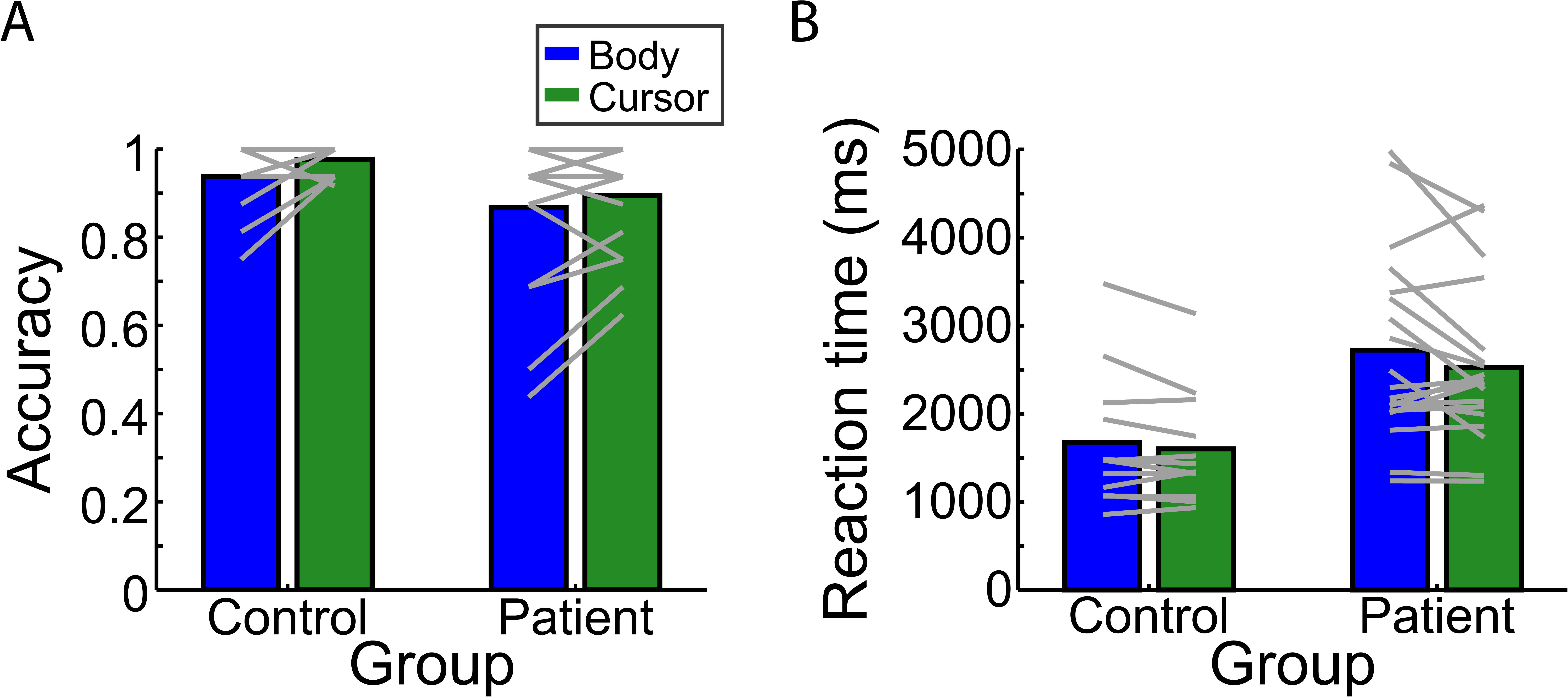
Mean performance in the Perception task was quantified using the complimentary measures of accuracy (A) and RT (B). Group means (bars) and individual means (lines) are shown separately for the body-cued (blue) and cursor-cued (green) conditions, for both patients and controls.

For response accuracy (Figure 5A), a mixed-effects model revealed no significant main effects: while there was a small difference between groups (χ^2^(1) = 5.229) and between stimulus types (body-cued versus cursor-cued: χ^2^(1) = 3.951), neither effect was significant after correcting for multiple comparisons (*p* = 0.11 and *p* = 0.19 respectively). No significant interaction was observed between group and stimulus type (χ^2^(1) = 0.445, *p* = 0.45).

For RT (Figure 5B), a mixed-effects model showed a significant main effect of group χ^2^(2) = 14.89, *p* = 0.0023), with patients having longer RTs than controls (2623.7 ± 948.3 ms and 1638.4 ± 696.0 ms respectively). We observed no effect of testing order (χ^2^(1) = 2.49, *p* = 0.114) nor any interaction between order and group or stimulus type. Importantly, we again observed no significant effect of stimulus type (χ^2^(1) = 2.633, *p* = 0.209) and no interaction between stimulus type and group (χ^2^(1) = 0.450, *p* = 0.502). Together, the accuracy and RT data suggest that patients in general exhibited disrupted performance on the Perception task compared to that of control participants, and performed similarly when the movement was cued by an actor or a cursor. Thus, as with the Production task, availability of information about the relative positioning of the limb segments did not strongly influence the ability to accurately perceive and report the observed trajectory path.

### Performance on the Production and Perception tasks were uncorrelated

The previous two analyses indicated that left hemisphere damage disrupts performance on both the Production and Perception tasks. Since these analyses were performed at the group level, it is important to understand whether the same participants exhibited deficits on both tasks (indicating that both tasks potentially rely on the same underlying computational processes or that difficulties in the Perception task can account for impaired performance in the Production task), or whether the group-level deficits stemmed from different subgroups of patients that were impaired on only one of the two tasks (suggesting that separate processes may underlie performance of the Production and Perception tasks).

The relationships between shape dissimilarity in the Production task and performance (accuracy and RT) in the Perception task are summarized in Figure 6. This figure suggests that there is no clear relationship between performance on the two tasks. In fact, several patients exhibited relatively good performance in the Production task (low shape dissimilarity) but disrupted performance on the Perception task (low accuracy/high RT), while other patients exhibited the opposite trend.

**Figure 6.**
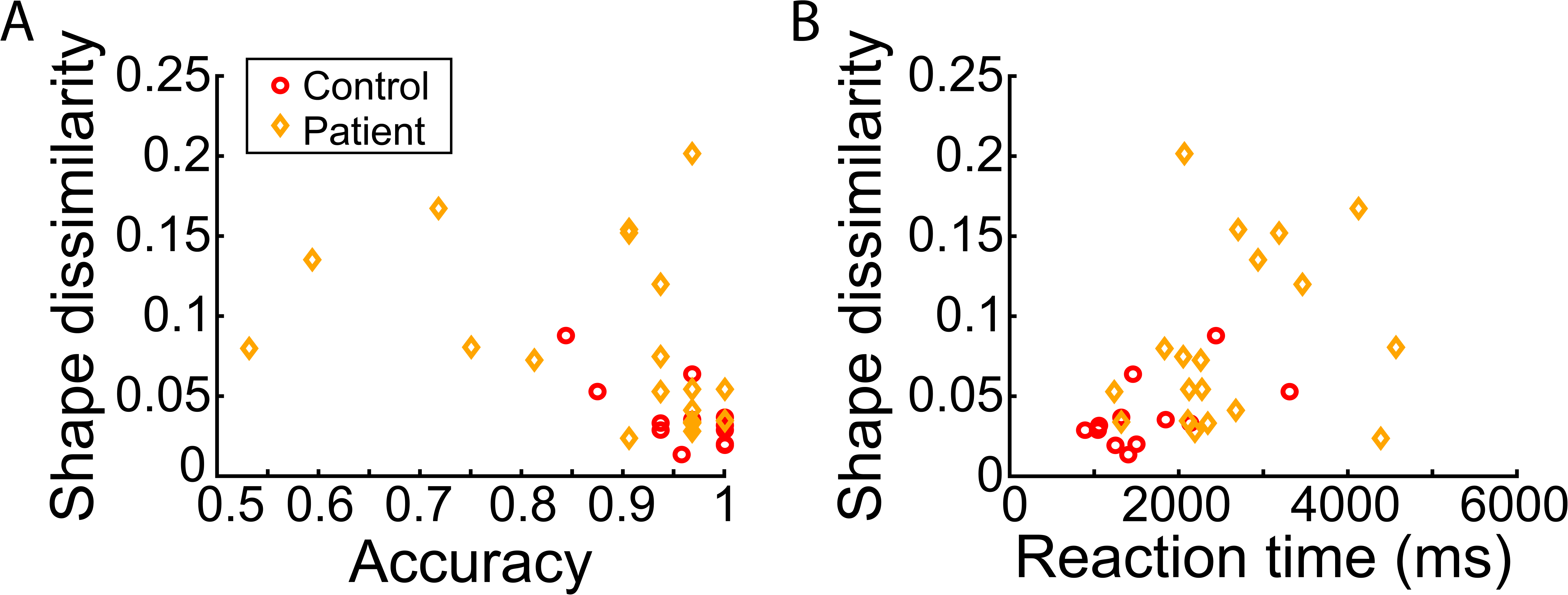
Relationship between Production (ordinate) and Perception (abscissa) tasks, shown separately for Perception task accuracy (A) and RT (B). Mean performance is shown for each patient (orange diamonds) and control (red circles).

To confirm these impressions and to guard against outliers more robustly, we used mixed-effects models to predict mean Production-task shape dissimilarity using either mean Perception-task accuracy or mean Perception-task RT (see Methods). These models also accounted for the effects of primary task factors (e.g., group and stimulus type) and a random effect of participant, and in the case of RT also accounted for performance on the Memory control task. In this analysis, we found no significant effect of Perception task accuracy (χ^2^(1) = 1.07, *p* = 0.30) or RT (χ^2^(1) = 0.070, *p* = 0.79) on shape dissimilarity, suggesting that how well a given individual performed on the Production task was uncorrelated with their performance during the Perception task. This was also true when looking only at patients (effect of accuracy: χ^2^(1) = 0.028, *p* = 0.37; effect of RT: χ^2^(2) = 0.24, *p* = 0.62) or only at controls (effect of accuracy: χ^2^(2) = 0.027, *p* = 0.87; effect of RT: χ^2^(2) = 2.37, *p* = 0.12). Thus, across both groups we observed a dissociation between how participants performed on the Perception and Production tasks, which indicates that a subset of participants had difficulties specifically in the ability to imitate but not to simply report the shape of the desired movement.

### Lesion analyses

Given the heterogeneity of patient performance in both the Production and Perception tasks, our final set of analyses focused on understanding this variability based on lesion location. We utilized two complementary lesion-analysis approaches. First, we performed Support Vector Regression-Lesion Symptom Mapping (SVR-LSM) analyses (Zhang et al., 2014) to examine the voxel-level relationship between performance in the behavioral tasks (shape dissimilarity for the Production task and both the accuracy and RT measures for the Perception task) and the patients’ lesion locations. Second, we performed a Region of Interest (ROI) analysis (Buxbaum et al., 2014) to examine the contributions of specific brain areas (defined on the lesion maps according to Brodmann areas) that we hypothesized to be involved in imitation based on prior research; this included PMd (approximated using Brodmann area 6, although we recognize that Brodmann area 6 also includes PMv) (Hocherman and Wise, 1990; Caspers et al., 2010; Pearce and Moran, 2012), and IPL (Brodmann areas 39 and 40) (Buxbaum et al., 2014; Kadmon Harpaz et al., 2014). The ROI analyses allowed us to identify correlations between damage to larger neural *regions* and behavioral impairments – effects that might be missed with a voxel-level analysis like SVR-LSM, particularly if there is inter-subject variability in the individual voxels associated with the task. Importantly, the results of both lesion analysis approaches were found to be consistent; we report the outcomes of both approaches below.

The SVR-LSM analysis of lesions associated with shape dissimilarity in the Production task (averaging across stimulus type) is shown in Figure 7A. The lesions contributing most highly to increased shape dissimilarity were located in the dorsal premotor and primary motor cortices. These regions were highly overlapping for the body and cursor conditions when analyzed separately (Figure 7B), although there was some suggestion that regions associated with poor performance in the body-cued condition extended more ventrally while those in the cursor-cued condition extended more dorsally.

**Figure 7.**
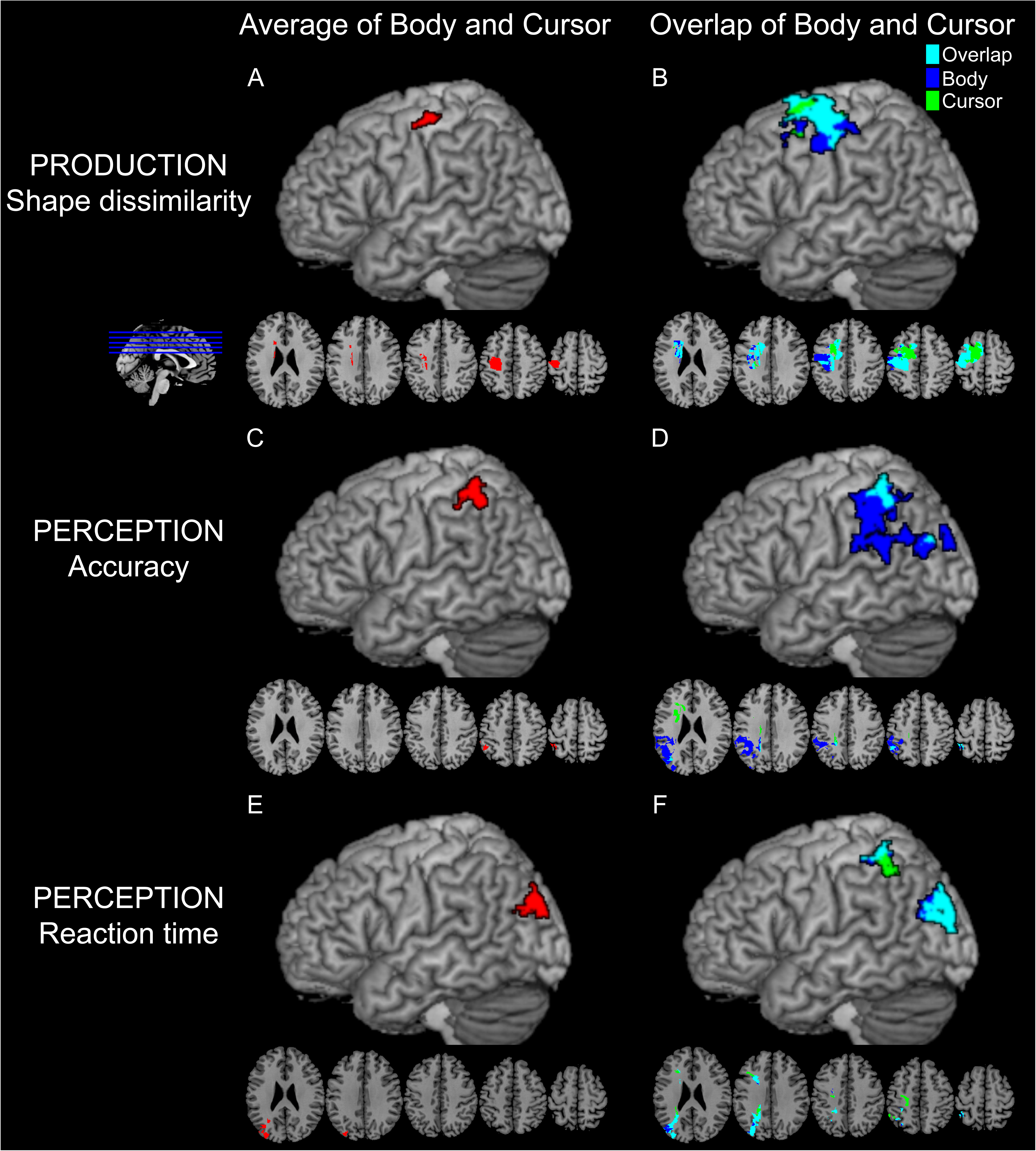
Results of the SVR-LSM analyses (N = 21) of shape dissimilarity in the Production task (top row), Perception task accuracy (middle row), and Perception task RT (bottom row). Results are presented for both the average data collapsing across the body and cursor stimuli (left column), as well as displaying the body and cursor results separately as well as their overlap at a relaxed statistical threshold (right column).

In contrast, the SVR-LSM analyses for the Perception task suggested that accuracy in reporting the movement shape is associated with damage to the supramarginal gyrus (Figures 7C). Interestingly, while there was a high degree of overlap of lesions associated with worse accuracy in the body-cued and cursor-cued conditions separately (Fig. 7D), areas associated with worse performance in the body-cued condition were much larger and extended ventrally to encompass areas traditionally associated with the ventro-dorsal stream and imitation ability (Buxbaum et al., 2014). In contrast, Perception task RT showed a slightly different pattern, being predominantly associated with damage to middle occipital gyrus (Fig. 7E). There was also a high degree of overlap in lesions associated with slowed RTs when considering body-cued and cursor-cued conditions separately (Fig. 7F), primarily in middle occipital gyrus but also in the middle intraparietal sulcus.

Thus in general, the findings from the SVR-LSM analysis suggest that lesions to different regions of the brain were associated with performance in the Production and Perception tasks respectively. The Production task was largely correlated with lesions in PMd and M1, while the Perception task was associated with lesions in the IPL. Furthermore, in both tasks, lesions that were associated with behavioral deficits due to cuing using a body-based or body-free stimulus substantially overlapped. However, there was a tendency for the regions associated with deficient performance with body stimuli to be more inferior than those associated with the overlap of body and cursor conditions, particularly in terms of accuracy performing the Perception task.

The SVR-LSM analysis looks for consistently lesioned continuous groups of voxels across all subjects that are associated with behavioral impairments. However, because of our relatively small sample size and the fact that stroke lesion sites do not follow predefined brain regions, we verified our findings using a ROI analysis approach using regions selected *a priori* that included PMd (Brodmann area 6, PM) and the IPL (Brodmann areas 39 and 40). In doing so, we found results consistent with the SVR-LSM analysis. Specifically, for the Production task, the proportion of damage to Brodmann area 6 (premotor cortex) significantly predicted shape dissimilarity (χ^2^(1) = 6.22, *p* = 0.050). In contrast, for the Perception task, there was a significant correlation between accuracy and proportion damage to Brodmann area 40 (supramarginal gyrus) (χ^2^(1) = 6.67, *p* = 0.039). We did not observe any other correlations between these regions and the performance metrics from the Production or Perception tasks, and there was no significant effect of stimulus type in any cases.

Taken together, the results of the two lesion-analysis approaches, along with the lack of correlation in performance across patients for the Production and Perception tasks, suggest a dissociation between the brain regions required to perform the Production and Perception tasks.

## DISCUSSION

Although imitation is an important aspect of learning, communicating, and using tools, the mechanisms by which people imitate are poorly understood. Here we demonstrated that individuals can imitate meaningless actions by representing the trajectory of the end-effector’s movement through space, and that this ability is disrupted by a stroke affecting the left PMd: cueing trajectories by an actor or a cursor yield similar performance impairments in patients. Such impairments cannot be attributed to general deficits due to a stroke, since they were observed in the non-paretic arm and patients had no difficulty imitating point-to-point gestures. Moreover, imitation deficits do not reflect visual working memory deficits. Together, these findings support the existence of a body-independent, trajectory-based route for imitation. Finally, behavioral and anatomic dissociations between the imitation of movement trajectories and the ability to identify their shapes suggest distinct roles for parietal and premotor cortices in imitation.

### Imitation by representing the trajectory path

In our task, patients and controls performed similarly when imitating trajectories cued by an actor (providing information about whole-limb configurations) or a cursor (providing information only about motion of the end-effector). This suggests that gestures can be imitated by representing only the trajectory path of the movement, consistent with research suggesting that participants attend only to the portion of the arm most relevant for imitation (Matarića and Pomplunb, 1998). Interestingly, a body-independent trajectory-based route for imitation could explain how species such as dolphins or parrots imitate humans (Moore, 1992; Harley et al., 1998; Xitco Jr et al., 1998) despite lacking homologous limbs.

Prior research has suggested the importance of planning end-effector trajectories (Abend et al., 1982; Flash and Hogan, 1985). Indeed, hand position variance is smaller than that of the elbow or shoulder (Sergio and Scott, 1998; Tseng et al., 2002). Our previous research suggests that neurotypical individuals plan curved trajectory paths when reaching around obstacles (Wong et al., 2016), and can guide corrective responses to unexpected perturbations (Cluff and Scott, 2015). Finally, trajectory representations can explain the kinematic invariance of letters written with different effectors (Wright, 1990; Rijntjes et al., 1999) or at different sizes (van Galen and Teulings, 1983; Wing, 2000).

Gestures could be represented compactly by only specifying movement trajectories. Rather than recalling the motion of every limb joint, only the path of the end-effector must be recalled; hence, describing a gesture becomes more computationally tractable (Ijspeert et al., 2001; Billard et al., 2004). Furthermore, because trajectory representations are not directly tied to the details of movement execution (e.g., controlling particular muscles), a single representation can support movement with either arm; this provides one potential explanation for how a unilateral lesion in the left hemisphere could disrupt movement imitation bilaterally.

Finally, our data indicate that PMd is associated with the ability to plan trajectory paths. This region has previously been associated with the dorso-dorsal stream (Rizzolatti and Matelli, 2003; Binkofski and Buxbaum, 2013), which is primarily associated with planning visually-guided reaches. However, the dorso-dorsal stream has also been associated with imitation (Rumiati et al., 2005; Caspers et al., 2010; Hoeren et al., 2014), and neural activity in PMd modulates with hand trajectory (Hocherman and Wise, 1990, 1991; Pearce and Moran, 2012; Pilacinski et al., 2018). Our findings thus suggest that PMd, and the dorso-dorsal stream in general, is important for planning actions when the goal can be specified in a body-independent manner (e.g., a location in space or trajectory path).

Planning trajectory paths could be more broadly useful for praxis (imitating gestures, pantomiming, or using tools). Tool use, for example, could be guided by planning the tool’s trajectory rather than (or in addition to) the potentially incongruous movement of one’s body. Such a trajectory representation is consistent with previous descriptions positing that imitation and tool-use abilities depend upon representations of “time-space-form pictures” of movements (Liepmann, 1905; Geschwind, 1975; Mack et al., 1993).

### Dissociation between production and perceptual reporting

In addition to trajectory-imitation deficits, stroke participants also had difficulties reporting the shapes of observed trajectories. Curiously, poor performance in producing trajectory paths was not correlated with difficulty in reporting trajectory shapes. Moreover, these tasks dissociated neuroanatomically: deficits in the Production task were associated with lesions in PMd (part of Brodmann area 6), whereas behavioral deficits in the Perception task were associated with lesions in the IPL (supramarginal gyrus, Brodmann area 40) even after controlling for potential visual working memory impairments. Moreover, we observed a greater dissociation between body-cued and cursor-cued performance in the Perception task, with lesions associated with performance in the body condition being more ventral; this suggests that while production of movement trajectories may be similar regardless of the input route, earlier processing is more distinct for body-based and cursor-based stimuli.

Although it seems surprising that participants could be impaired only in the Perception task, this could arise for a couple of reasons. For one, the Perception task requires recognizing the similarity between the stimulus motion and the static shape, whereas the Production task requires recall of the action. A lack of correlation between the Production and Perception tasks is therefore consistent with the distinction between recognition and recall (Hollingworth, 1913; Kintsch, 1970; Anderson and Bower, 1972; Tulving, 1976). Additionally, the Perception task may uniquely require the transformation from a dynamic movement to a static image (Korneev and Kurgansky, 2014), or the ability to mentally rotate body parts (Bonda et al., 1995; Zacks, 2008) and distinguish mirror-reversed shapes (Davidoff and Warrington, 2001) which are both associated with neural activity in parietal cortex. Thus, behavioral dissociations may arise not from perception of the stimulus motion per se but from secondary processes that are not shared between the two tasks.

### Two routes to imitation

Although it is possible to imitate end-effector trajectories, under certain circumstances it is also important to copy the positioning of the entire limb. For example, a coach may instruct an athlete to modify her elbow position while swinging a tennis racket. Planning trajectories seems poorly suited for this, as it would require a trajectory representation for both the racket and the elbow (Ijspeert et al., 2001), and it has been shown that planning multiple trajectories simultaneously is challenging (Albert and Ivry, 2009). In such situations, representing limb configurations would be useful for imitation, e.g., by specifying the transformations required to reconfigure the limb (Chaminade et al., 2005; Amorim et al., 2006). Thus, imitation might be carried out in two ways: by copying a movement trajectory or a body-part-relationship goal. Indeed, kinematic errors can dissociate from postural-configuration errors in imitation (Hermsdorfer et al., 1996).

Claims regarding a body-dependent route to imitation are based on studies in which imitation ability is scored according to the successful positioning of body segments (elbow, hand, etc.) (Tarhan et al., 2015), often with respect to a static goal posture (Goldenberg, 1995; Goldenberg and Hagmann, 1997; Harrington and Haaland, 1997). Thus it is not surprising that these studies support a body-specific representation for imitation (Goldenberg, 1999; Buxbaum et al., 2000; Schwoebel et al., 2004; Goldenberg, 2013). In contrast, we emphasized trajectory imitation and did not assess body-configuration errors. Regardless, body-based representations seem most useful when imitating static postures. Imitating dynamic gestures would require an additional step, such as linking sequences of static body configurations. In contrast, trajectory-based representations seem more amenable for imitating dynamic gestures as they describe spatiotemporal information about motion along the desired path (e.g., instantaneous velocity).

Previously proposed neuroanatomical models of action planning have divided the dorsal stream (Ungerleider and Mishkin, 1982; Milner and Goodale, 1995) into two routes for action (Rizzolatti and Matelli, 2003; Binkofski and Buxbaum, 2013). Body-based representations have been associated with the ventro-dorsal pathway passing through the IPL (Goldenberg, 1995, 1999; Buxbaum et al., 2000; Schwoebel et al., 2004). In contrast, the dorso-dorsal stream has been associated with planning movements to targets in extrinsic space, and may thus represent actions in a body-independent manner. Our finding of a body-independent trajectory representation for imitation in PMd is consistent with the role of this latter pathway. Interestingly, our data also suggest that performance on body-cued trials may be associated with more ventral regions than that of cursor-cued trials, consistent with the dual-route model. Nevertheless, the existence of overlapping regions associated with body-cued versus cursor-cued performance in both the Production and Perception tasks suggest the existence of a route from perception to action specifically concerned with representing movement trajectories.

### Conclusions

This study tested the hypothesis, based on our prior research, that humans possess an ability to represent movement trajectories that is useful for imitating meaningless actions. It also sought to identify where in the praxis network spanning the left fronto-parietal cortex such a trajectory-path representation exists. Our findings confirm the existence of an abstract (body-independent) trajectory-path representation for planning and executing complex actions such as copying gestures, which can be dissociated behaviorally and neuroanatomically from the ability to accurately report the shape of the observed action. Furthermore, we provide strong evidence that damage to dorsal premotor cortex disrupts this ability to represent trajectory shapes, giving rise to deficits in the ability to imitate meaningless actions.

## AUTHOR CONTRIBUTIONS

S.A.J. and A.L.W. conceived and performed experiments, analyzed the data, and wrote and edited the manuscript. L.L.S. performed the experiments and processed the data for analysis. L.J.B. and J.W.K. conceived the experiments, supervised the project, edited the manuscript, and provided resources and funding.

## ACKNOWLEDGEMENTS

This work was supported by NSF Grant BCS-1358756 to JWK and NIH Grant R01-NS065049 to LJB. The authors would like to thank H. Branch Coslett for his assistance with lesion identification, Stephanie Shaia and Harrison Stoll for their assistance with the SVR-LSM analyses, and Peter Turkeltaub and Andrew DeMarco for sharing their SVR-LSM software. The authors declare no competing financial interests.

## Footnotes

1 Participants also saw a third ‘stick’ stimulus-type was intended to test whether individuals would imitate similarly in response to an actor or to jointed non-body stimuli. The location of the actor’s shoulder, elbow, and fingertip locations were used to produce a two-segment stick-figure arm moving in 2D. However, all participants found this condition confusing due to projection of the 3D arm motion into a 2D plane, as confirmed by reliably worse performance compared to the other conditions across all participants and tasks. Since this condition did not offer meaningful insight into trajectory imitation per se, it was excluded from further analysis, but we mention it here for completeness as the trials of all three stimulus types were interleaved throughout the task.

